# Detection of Aggressive Mesenchymal Glioblastoma by Mannose-Weighted CEST MRI

**DOI:** 10.1101/2025.04.15.648948

**Authors:** Behnaz Ghaemi, Hernando Lopez-Bertoni, Shreyas Kuddannaya, Sophie Sall, John Laterra, Guanshu Liu, Jeff W.M. Bulte

## Abstract

Glioblastoma (GBM) contain mesenchymal cancer stem cells that drive tumor aggressiveness and recurrence and exhibit aberrant glycosylation during proneural-to-mesenchymal transition. A comprehensive analysis of human GBM transcriptomic datasets revealed an upregulation of 13 genes involved in mannosylation. Histopathological staining of a tissue array representing 35 GBM cases revealed elevated mannose, correlating with increased expression of the mesenchymal marker CD44. Mannose-weighted chemical exchange saturation transfer magnetic resonance imaging (MANw CEST MRI) detected elevated mannose levels in aggressive mesenchymal GBM neurospheres *in vitro* and *in vivo*, but not in less aggressive non-mesenchymal phenotype. To establish causation, inhibiting the expression of the mannose binding lectins LMAN1/2 that regulate intracellular processing of mannosylated proteins decreased the glioma cell MANw CEST MRI signal. Our findings indicate that MANw CEST MRI can visualize high mannose levels in mesenchymal GBM cells, which may serve as a surrogate imaging biomarker for predicting and assessing tumor aggressiveness and recurrence.

---

Glioblastoma (GBM) is a very aggressive primary brain tumor with poor prognosis and high incidence of recurrence^1,2^. Gene expression profiling studies in GBM have revealed distinct molecular subtypes (e.g . classical, proneural, and mesenchymal) associated with unique genomic abnormalities and clinical outcomes^3^. The mesenchymal^3^ and mesenchymal-like^4^ subtypes associate with the most aggressive disease, worse prognosis, resistance to current therapies and recurrence^5^. Identifying key features of this mesenchymal transition can allow for the development of novel biomarkers to enable early detection of aggressive cell subset and improve patient outcome.

Protein glycosylation plays a critical role in multiple cellular processes including cell signaling, cell-extracellular matrix interactions, cell migration, immune modulation, and the maintenance of tissue structure and homeostasis^6^. Among these, mannose-containing glycoproteins stand out due to their involvement in cellular recognition, trafficking, and immune response regulation^7^. Aberrant glycosylation, including alterations in mannose residues, occurs during the epithelial-to-mesenchymal transition (EMT) and can enhance cell motility, invasiveness, and survival of tumor cells^8^. Understanding glycoprotein biology, particularly the role of mannose-enriched glycans in tumor cells and their microenvironment, can allow for identification of novel biomarkers and therapeutic strategies to positively impact cancer patient outcome.

Currently, precision imaging in oncology has limitations, as dictated by the heterogeneity and spatiotemporal composition of tumor cells and their microenvironment^9^. In this regard, advancing molecular imaging may allow us to leverage aberrant glycan biology for diagnostic purposes. Chemical exchange saturation transfer magnetic resonance imaging (CEST MRI) is an emerging molecular imaging technique that can detect metabolites and molecules that contain exchangeable amide, amine, and hydroxyl protons^10,11^. The hydroxyl (OH) protons present in sugar residues have been exploited as an endogenous CEST MRI biomarker for mucin underglycosylation in adenocarcinoma^12^ and as exogenous glucose^13–16^ and dextran^17,18^ CEST MRI contrast agents for imaging tumor perfusion and blood-brain barrier leakage. We have recently shown that human mesenchymal stem cells (hMSCs) can be detected by mannose-weighted (MANw) CEST MRI by virtue of the high content of their cell membrane mannose N-linked glycans^19^. These findings prompted us to investigate if MANw CEST MRI can be used to differentiate aggressive GBM with a mesenchymal phenotype from less aggressive non-mesenchymal GBM, with the ultimate goal to develop the MANw CEST MRI signal as a surrogate imaging biomarker for tumor aggressiveness, recurrence and subtype classification in patients.

## RESULTS

### Mesenchymal GBM cells express high levels of mannose and genes regulating mannosylation

Since mesenchymal transitions play important roles in GBM pathogenesis^3–5,20^, we aimed to identify molecular events related to these transitions that occur concurrently with changes in mannosylation. We queried expression of known genes involved in the generation and processing of mannosylated glycans in 3 distinct clinical GBM transcriptomic data sets. This analysis identified 13 relevant genes (LMAN1, LMAN2, LMAN2L, SLC35D2, TMEM5 DPY19L1, DPM3, DPM2, DPM1, DPY19L4, PIGB, FKTN, POMT2) as being consistently upregulated in GBM compared to normal brain **(Fig. 1A)**. Further analysis showed that expression of 4 of these 13 genes (LMAN1, LMAN2, LMAN2L and SLC35D2) is elevated in mesenchymal GBM cells compared to the proneural subtype **(Fig. 1B)**. We also identified positive correlations between LMAN1, LMAN2, SLC35D2 and the mesenchymal marker CD44, but not the proneural marker Prom1 (CD133) in transcriptomic datasets derived from clinical specimens **(Fig. 1C)** and GBM stem cells **(Fig. 1D).** Consistent with these transcriptomic associations, fluorescence-activated cell sorting (FACS) followed by quantitative reverse transcription polymerase chain reaction (qRT-PCR) determined that LMAN1 and LMAN2 are enriched in CD44^+^ GBM cells compared to their CD44^-^ counterparts (**Fig. 1E**).

**Figure 1:**
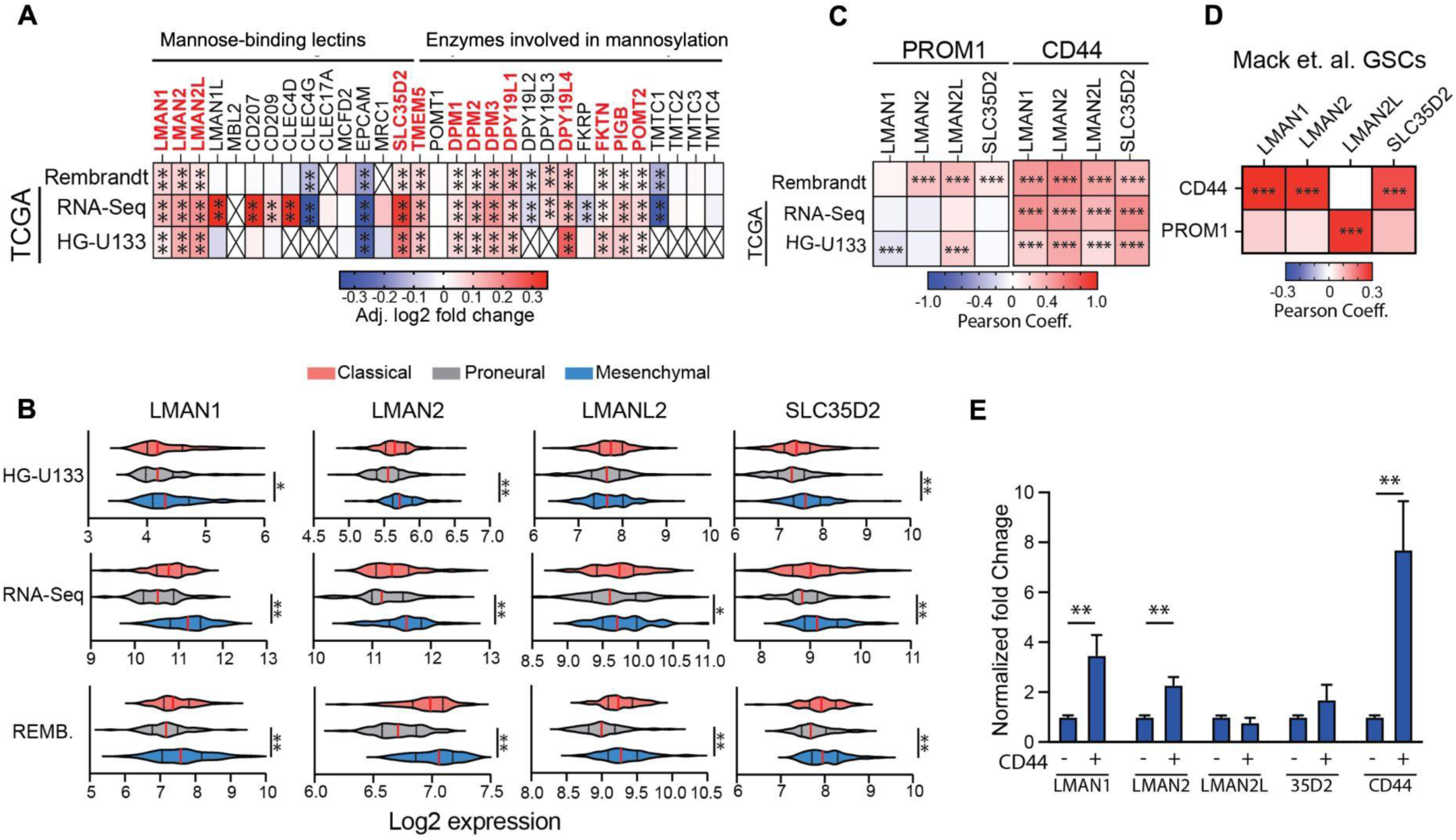
LMAN1 and LMAN2 are enriched in mesenchymal GBM cells. **(A)** Expression of genes coding for regulators of mannose metabolism was compared between IDH-wt GBM and normal brain tissue. Upregulated genes are shown in red. **(B)** Violin plots showing expression of LMAN1, LMAN2, LMAN2L, and SLC35D2 in different GBM molecular subtypes. Correlation matrix showing Pearson’s coefficients between the mesenchymal marker CD44 or prominin and LMAN1, LMAN2, LMAN2L, and SLC35D2 in GBM clinical specimens **(C)** and primary GBM neurospheres **(D)**. **(E)** qRT-PCR to measure expression of LMAN1, LMAN2, LMAN2L, and SLC35D2 in GBM neurospheres sorted into CD44 positive and negative populations. Statistical significance was calculated using unpaired, non-parametric, student’s t-test with Mann-Whitney post hoc test in panels **A, B;** statistical significance was calculated using Student’s t-test in panel **E.** Expression data was retrieved from the clinical databases using the GlioVis portal (http://gliovis.bioinfo.cnio.es). *p<0.05, **p<0.01; ***p<0.001; X=not determined.

To compare glycan mannosylation and CD44 expression between GBM and normal brain in patient samples, a tissue array consisting of 35 glioblastoma and 5 normal brain specimens was analyzed. Fluorescence imaging revealed an association between mannose and expression of mesenchymal markers CD44 in GBM samples (**Fig. 2A,B**), whereas normal brain tissue exhibited little to no expression of these markers. Analysis of CD44 expression and mannose levels demonstrated a positive correlation (r=0.65, p=0.0003)(**Fig. 2C**), indicating a connection between elevated mannose levels and mesenchymal phenotypic transitions in GBM.

**Figure 2:**
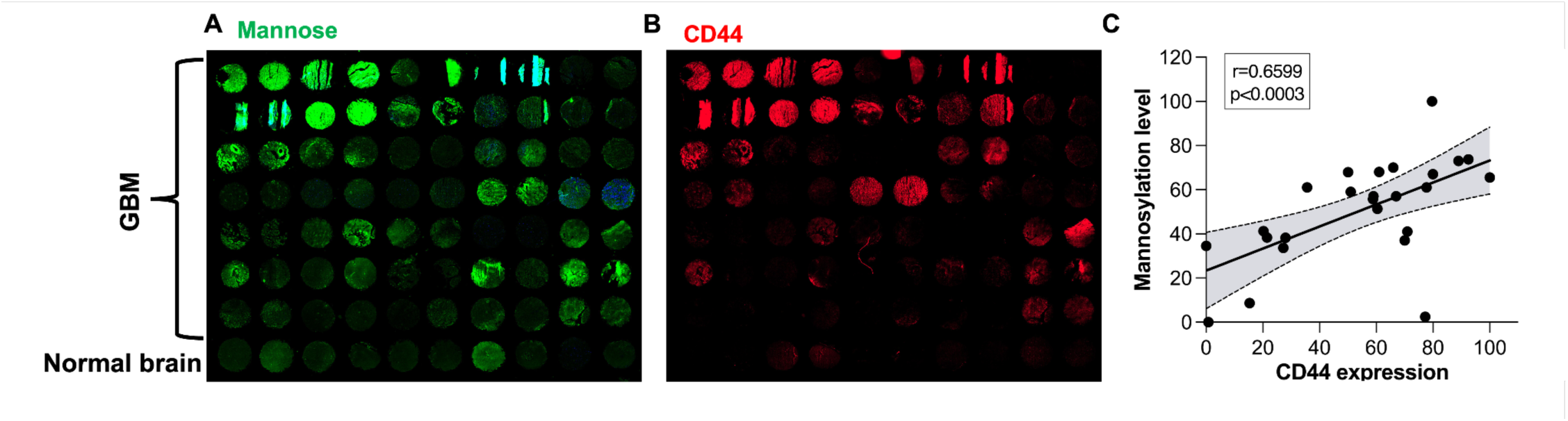
Tissue array of GBM reveals the co-presence of mannose and CD44, which correlate to tumor aggressiveness. Clinical tissue array containing 35 different GBM (patients aged 8-80 years) and 5 normal human cerebral tissue samples (for a total of 80 in duplicate) were stained for (**A**) mannose and (**B**) the mesenchymal marker CD44. (**C**) Scatter plots with linear regression lines show the relationship between CD44 expression and mannose levels in GBM tissue samples. Shaded gray areas represent 95% confidence intervals.

### *In vitro* MANw CEST MRI of proneural and mesenchymal GBM correlates to mannose and CD44 co-expression

Mesenchymal transitions and changes in mannose levels are critical determinants of the tumor and stem cell phenotype in multiple cancers, including GBM^8,21,22^. Our findings so far support the hypothesis that CD44+ mesenchymal GBM cells express high levels of mannose compared to their proneural counterparts and that these molecular differences can be leveraged for label-free MANw CEST MRI detection **(Fig. 3A)**. To test this hypothesis, we used two well-characterized patient-derived human isocitrate dehydrogenase (IDH)-wt GBM neurosphere cell lines that represent the mesenchymal (M1123) and proneural subtypes (GBM1a)^23,24^. As expected, GBM1a expresses high levels of the proneural markers CD133, Sox2, and Ascl1 while M1123 expresses high levels of the mesenchymal markers CD44, vimentin (Vim) and Snai1 **(Fig. 3B,C)**. Consistent with these distinct molecular features of proneural and mesenchymal GBM cells, M1123 cells also expressed higher amounts of mannose when cultured as neurospheres in defined serum-free stem cell medium, but not when cultured under differentiating conditions in serum-containing medium **(Fig. 3D, S1A).** Accordingly, the higher levels of mannose correspond to a significantly (p<0.05) higher MANw CEST signal at 1.2 ppm, i.e., the CEST signature of OH protons abundantly present in mannose (**Fig. 3E**). Importantly, we observed a significant decrease in mannosylation (**Fig. S2A**) and MANw CEST signal (**Fig. 3F and S1B-D**) in M1123 neurospheres upon serum-induced cell differentiation, conditions that reduce the stem cell phenotype of GBM cells^25,26^.

**Figure 3:**
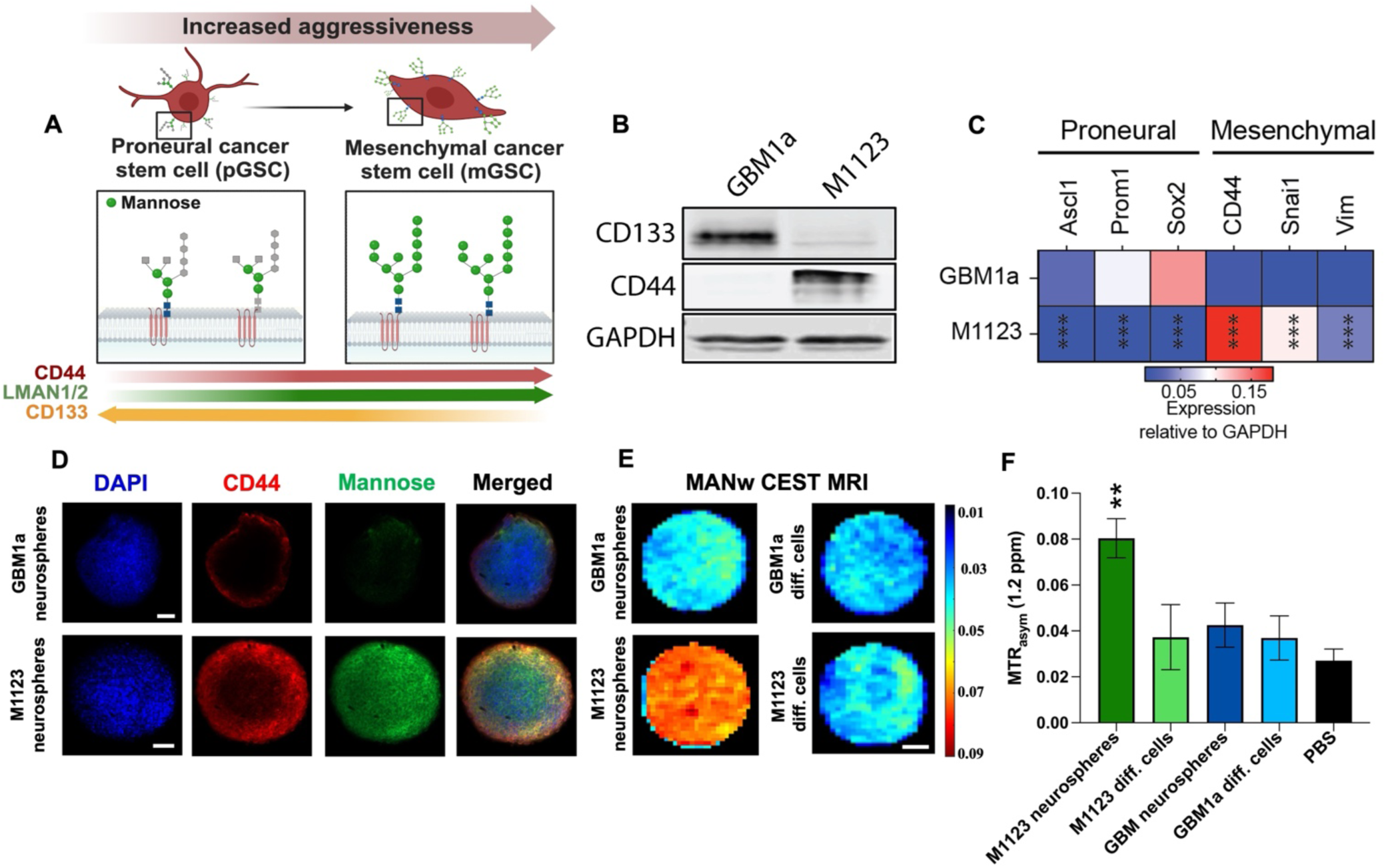
*In vitro* MANw CEST MRI signal corresponds to mannose and CD44 co-expression in mesenchymal GBM which is absent in proneural GBM. (**A**) Schematic illustration of the proneural-to-mesenchymal transition in GBM, resulting in elevated mannose N-linked glycan levels in CD44^+^ mesenchymal cells which is absent in CD133^+^ proneural cells. (**B**) Western blot showing expression of CD133 (proneural) and CD144 (mesenchymal) markers in GBM1a and M1123 cells. (**C**) qRT-PCR analysis of canonical proneural and mesenchymal markers in GBM1a and M1123 cells. **(D)** Mannose and CD44 staining in single representative GBM1a and M1123 neurospheres. Scale bar=200 µm. (**E**) *In vitro* MANw CEST MRI and (**F**) signal quantification of GBM1a and M1123 neurospheres suspended in a 5 mm NMR tube. Scale bar in (**E**)=1 mm. Statistical significance was determined using Student’s t-test, where **p<0.01.

### Endogenous mannose-binding lectins LMAN1 and LMAN2 impact mannose levels in mesenchymal GBM cells

Our computational analyses of GBM clinical specimens identified LMAN1 and LMAN2 as potential regulators of mannose homeostasis in mesenchymal GBM cells. Therefore, we examined the effects of LMAN1 and/or LMAN2 expression inhibition on mannose levels in M1123 neurosphere cells (**Fig. 4A**). qRT-PCR analysis demonstrated that the LMAN1- and LMAN2-specific siRNA oligos block their intended gene targets **(Fig. 4B)** concurrent with a significant reduction in mannose staining **(Fig. 4C,D)**. We did not observe an additive effect when we knocked-down LMAN1 and LMAN2 simultaneously, likely reflecting the known overlapping functions of these lectins in the intracellular processing of mannose glycans **(Fig. 4D)**. The decrease in mannose content after specific LMAN1 and LMAN2 knock-down correlated with a decrease in MANw CEST MRI signal *in vitro* **(Fig. 4E, Fig. S3)**.

**Figure 4:**
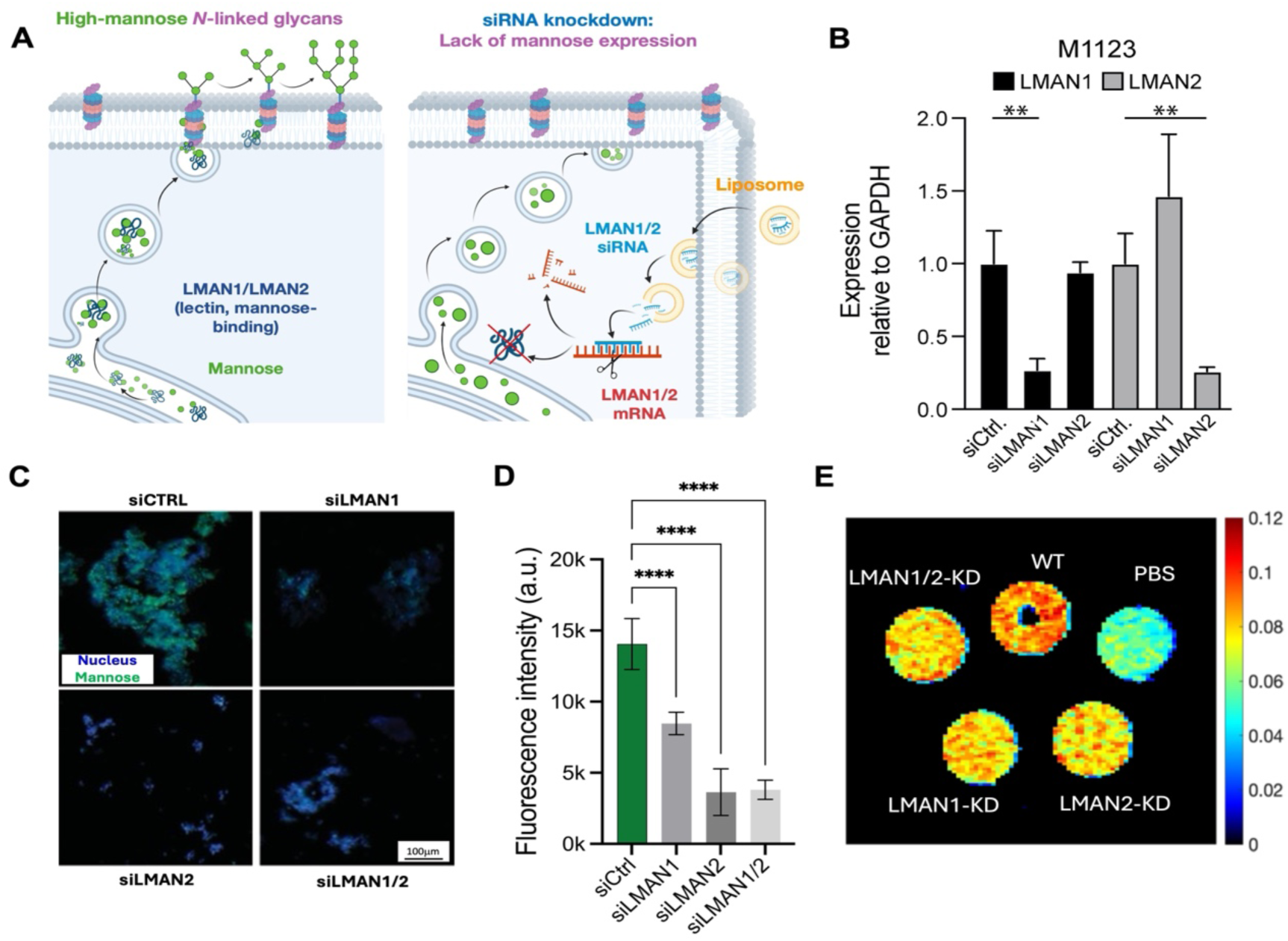
Knockdown of LMAN1 and LMAN2 in M1123 cells decreases mannosylation and MANw CEST contrast. (**A**) Schematic outline of siRNA-mediated silencing of LMAN1/2. (**B**) qRT PCR analysis shows specific knockdown of LMAN1 and LMAN2 compared to control siRNA and LMAN cross-control siRNA. (**C**) Mannose staining and (**D**) quantification. (**E**) *In vitro* MANw CEST MRI signal at 1.2 ppm for wild type (WT) cells and LMAN1/LMAN2 knockdown cells. Statistical significance was calculated using one-way ANOVA with Tukey’s post hoc test in panels B and D. Statistical significance was determined using one-way ANOVA, where **p<0.01 and ****p<0.0001.

### *In vivo* MANw CEST MRI of proneural and mesenchymal GBM correlates with mannose levels and CD44 expression

We assessed the feasibility of MANw CEST MRI in differentiating brain tumors derived from proneural vs. mesenchymal GBM neurospheres *in vivo*. Eight-week-old NOD SCID gamma mice received tumor cells in the left (GBM1a) and right (M1123) striatum. MRI was then performed on post-implantation days 1, 8, and 16. M1123-derived tumor xenografts exhibited a significantly higher growth rate concurrent with a prominent MANw CEST MRI signal that was absent in the GBM1a-derived tumors and normal brain. The MANw CEST signal in M1123 tumors remained consistently elevated (>1.8-fold) compared to GBM1a tumors and normal brain for all three time points, with a characteristic mannose peak around 1.2 ppm **(Fig. 5A,B, Fig. S3)**. We also analyzed tumors using APTw MRI^27^ a technique approved by the FDA in 2018 for its ability to identify active high-grade brain cancer by detecting endogenous proteins and peptides^28^. This is achieved by measuring the CEST signal at 3.6 ppm (amide protons) that is distinct from the 1.2 ppm signal from hydroxyl protons.

**Figure 5:**
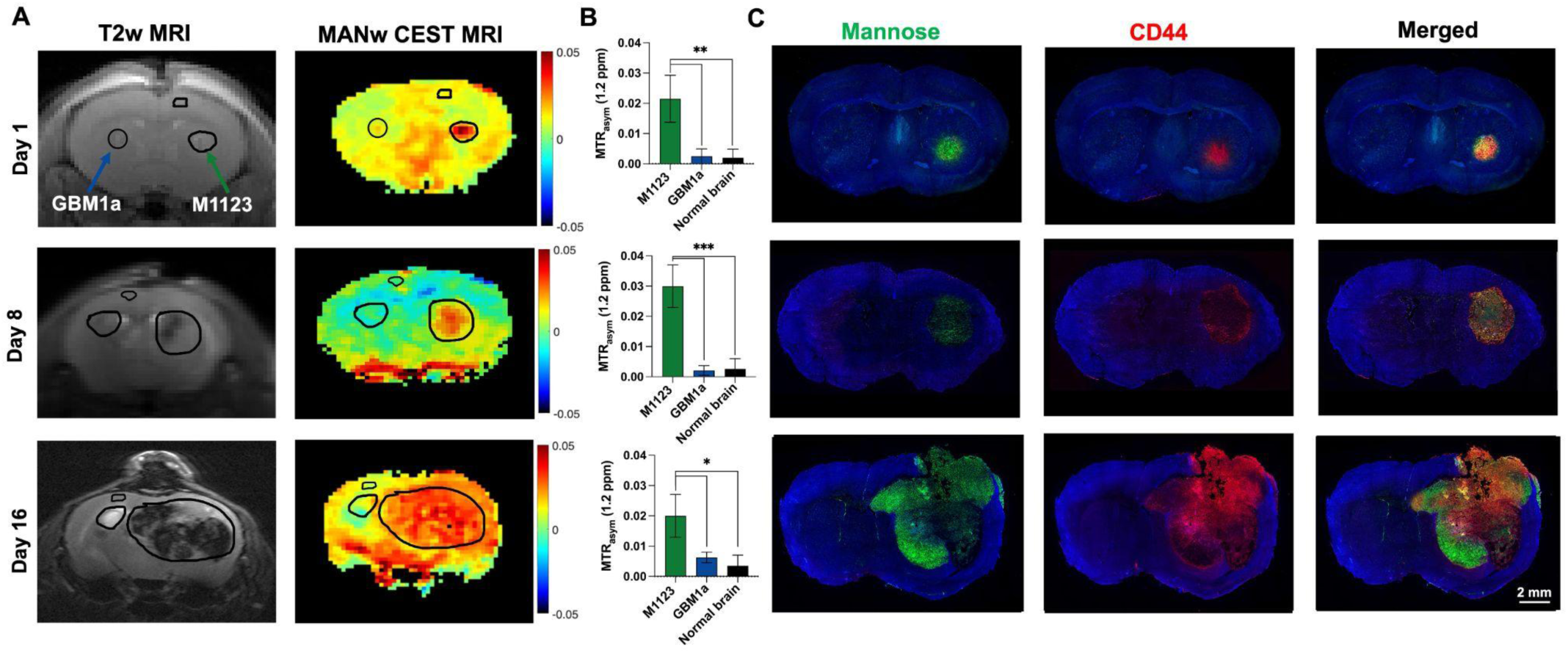
*In vivo* MANw CEST MRI signal corresponds to mannose and CD44 co-expression in mesenchymal GBM which is absent in proneural GBM. **(A)** Serial *in vivo* T2-weighted and MANw CEST MRI of a bilateral orthotopic GBM1a and M1123 mouse xenograft model at different time points post tumor implantation. (**B**) Quantified MANw CEST MRI signal at 1.2 ppm for the regions of interest drawn in A as an example for n=5 animals. (**C**) FITC-GNL and anti-CD44 staining in tissue sections representative for the brain slices shown in A. Statistical significance was determined using one-way ANOVA, where p<0.05; **p<0.01 and ***p<0.001.

Contrasting the MANw CEST MRI results, APTw MRI did not generate a consistent pattern in distinguishing GBM1a and M1123 tumors throughout the study (**Fig. S4**). APTw MRI signals from M1123 and GBM1a were indistinguishable on day 1. The M1123 APTw MRI signal was significantly higher than the GBM1a signal on day 8 and significantly lower than that from GBM1a signal on day 16 (**Fig. S4A,B**). Immunofluorescent post-mortem staining for mannose and CD44 colocalized with areas of MANw CEST MRI contrast, demonstrating that the cells with high mannose content represent mesenchymal GBM cells within the orthotopic M1123 tumor xenografts **(Fig. 5C)**. Gadolinium (Gd)-enhanced perfusion MRI, performed eight days post-tumor implantation, revealed no significant differences in perfusion characteristics between GBM1a and M1123 tumors, suggesting similar vascularization and blood-brain barrier permeability (**Fig. S5**).

GBM remains one of the most aggressive and treatment-resistant brain tumors, with 14-18 month median survival despite aggressive treatment^2,29^. Substantial efforts have been dedicated to characterizing the molecular landscape of GBM cells to understand mechanisms driving therapy resistance^4,30^. Gene expression profiling studies in GBM have highlighted mesenchymal transitions as a key contributor to poor patient outcomes^3,31^. Additionally, mesenchymal-like cancer cells exhibit enhanced motility, self-renewal capacity, and resistance to apoptosis and immune evasion^3,31^. Single-cell transcriptomic studies have further reinforced the link between mesenchymal transition and therapy resistance^5,32^. High-resolution single-cell transcriptomic analyses of GBM tumors before and after standard-of-care therapy demonstrated critical changes in the tumor microenvironment and cell state transitions associated with stemness-driving events and acquisition of mesenchymal signatures^32^.

Changes in glycosylation profiles are an important feature of mesenchymal transitions^8^ and offer a potential biomarker for classifying tumor subtypes and monitoring transitions to more aggressive treatment resistant tumors. Among various glycosylation patterns, high mannose-linked glycosylation is a distinct attribute of aggressiveness and may play a fundamental role in therapy resistance^33^. High-mannose glycosylation has been proposed as a biomarker for aggressive GBM subtypes^15^ and enhanced glycosylation is linked to hematopoetic stem cells^34^ and metabolic reprogramming of mesenchymal GBM stem cells^35^.

Interestingly, a recent study found that GSCs exhibit elevated levels of α-1,2-mannosidase (MAN1C1), an enzyme involved in the processing of N-glycans, and this elevated expression is associated with poor outcomes in glioma patients^36^. Changes in N-glycosylation driven by MGAT5, an enzyme that adds branching mannose residues to N-glycans, correlates with more invasive phenotype in GSCs^37^. Consistent with these observations, we show that mesenchymal GBM cells express higher levels of genes involved in mannose regulation **(Fig. 1)** and, for the first time, that mesenchymal GBM neurospheres display significantly higher mannose levels and higher MANw CEST signal than their proneural counterparts **(Figs. 3,4)**. Importantly, these correlations were maintained in patient derived glioma cells and tissues and tumor xenografts **(Fig. 1,2,4)**, underscoring the potential value of non-invasive imaging approaches to detect these molecular alterations in clinical settings.

While conventional T2-weighted and Gd-based contrast-enhanced MRI enable visualizing brain tumors, they lack specificity in differentiating tumor subtypes and assessing microenvironmental changes (**Fig. S5**)^38^. Advanced techniques such as diffusion-weighted imaging and perfusion imaging augment structural imaging with functional information by measuring water diffusion and vascularity^39^, respectively, while magnetic resonance spectroscopy further aids in tumor characterization by detecting metabolic markers such as choline and lactate^40^. However, these modalities insufficiently capture molecular features linked to specific tumor subtypes and their different aggressiveness. APTw MRI detects mobile proteins and peptides and has the ability to differentiate between high and low-grade glioma as well as to distinguish radiation necrosis from recurring tumors^41,42^. We now show that MANw is able to distinguish mesenchymal from proneural experimental GBM xenograft, which can be combined with APTw CEST MRI for imaging different and potentially complementary aspects of GBM pathology.

Like most cancers, GBM undergoes metabolic reprogramming to support tumor cell survival, proliferation, and invasion^43^. Aberrant glycosylation is a notable consequence of this metabolic rewiring, however, our understanding of the role of glycosylation in GBM pathology remains incomplete^44^. We have shown here that MANw CEST MRI can distinguish between proneural and mesenchymal cells and we rigorously show that this is based on their differential mannose content (**Fig. 3, 4**). Blocking expression of endogenous mannose-binding lectins, LMAN1 and LMAN2, significantly reduces MANw CEST MRI signal and mannose-specific lectin staining *in vitro* and *in vivo* **(Fig. 5)**, demonstrating that mannose is directly responsible as a contributor for MANw CEST MRI signal in mesenchymal GBM cells. As the MANw CEST MRI signal detects hydroxyl protons from all sugar residues, it is possible that mannose is not solely responsible for the increased signal, and in certain contexts other overexpressed glycans might also contribute. Integrating MANw CEST MRI with the currently available MRI portfolio, including APTw MRI, could enable a more comprehensive evaluation of GBM. We envision this approach will improve detection of aggressive mesenchymal tumors and their recurrence, refine treatment stratification, enhance the monitoring of therapeutic responses, and aid in optimizing surgical planning. This new imaging approach could be easily interfaced with current clinical imaging protocols to positively impact patient outcomes.

### Online content

Any methods, additional references, Nature Portfolio reporting summaries, source data, extended data, supplementary information, acknowledgements, peer review information; details of author contributions and competing interests; and statements of data and code availability are available at https://doi.org/xxx.

## Online Methods

### Patient databases

Clinical and transcriptomic data from control and glioma patient samples was retrieved from GlioVis database (http://gliovis.bioinfo.cnio.es/).

### Glioblastoma tissue array

A tissue microarray, containing 35 cases of glioblastoma and 5 cases of normal cerebral tissue (duplicate cores per case) were obtained from Tissuearray.Com (GL805-L51). CD44 expression and mannose levels were calculated by measuring mean fluorescence intensity (MFI) from fluorescence images, with values normalized on a 0–100% scale for comparison across samples.

### Cell culture

Human 0913 (GBM1a) cells were initially established by Vescovi and colleagues^23^ and characterized by our group^45^. M1123 cells were generously provided by Dr. Nakano (Ohio State University)^35^. Neurospheres were maintained in serum-free medium supplemented with epidermal growth factor (EGF) and fibroblast growth factor (FGF) and cultured in ultra-low-attachment 6-well plates (Millipore-Sigma) at 37°C under 5% CO₂/95% air. Serum-induced differentiation was performed by introducing serum into the culture medium and transitioning the cells to adherent conditions in T-25 flasks.

For the ELDA, tumor cells were cultured in stem cell medium containing EGF/FGF at decreasing cell densities ranging from 100 to 12.5 cells, with over 24 technical replicates for each dilution. The number of wells containing spheres was counted after 14 days, and the online ELDA tool (http://bioinf.wehi.edu.au/software/elda/, last accessed on 12.20.24) was used to calculate stem cell frequencies.

### Immunofluorescent staining for CD44 and mannose

GBM1a and M1123 neurospheres were transferred in 1-well cell culture chamber slides. Cells were first incubated in cold PBS containing 20 µg/ml GNL-FITC (FL-1241, Vector Laboratories) at 4 °C for 50 minutes for the presence of mannose, then washed three times with PBS. They were fixed with 4% paraformaldehyde for 20 min at room temperature (RT) and permeabilized using 0.1% Triton X-100 in PBS for 10 min. After blocking non-specific binding sites with 5% normal goat serum (NGS), cells were incubated with primary Alexa Fluor® 647 anti-CD44 antibody (1:250, Abcam, EPR18668) according to the manufacturer’s protocol (3 h at RT in the dark). Excess antibody was removed by washing 3x with 10 mM phosphate buffered saline, pH=7.4 (PBS). Nuclei were stained with 4′,6-diamidino-2-phenylindole (DAPI) for 5 minutes. To stain the tissue array slides, they were deparaffinized by sequential washes in xylene (2×5 minutes), followed by rehydration through a graded ethanol series (100%, 95%, 70%, and 50%, 5 minutes each) and a final rinse in distilled water. For antigen retrieval, slides were incubated in sodium citrate buffer (10 mM, pH=6.0) at 95–100°C for 20 minutes, then allowed to cool to RT before washing three times with PBS. For immunostaining, slides were blocked in 5% bovine serum albumin (BSA) in PBS for 1 hour at room temperature to minimize non-specific binding. Tissue arrays were then incubated with Alexa Fluor® 647 anti-CD44 antibody (1:250) and 20 µg/ml of GNL(Galanthus nivalis lectin)-FITC for 1 hour at RT in the dark. Unbound antibody/lectin was removed by washing three times with PBS. Nuclei were counterstained with DAPI for 5 minutes, followed by a final PBS wash. All fluorescence imaging was performed using a Zeiss Axiovert 200 M inverted epifluorescence microscope, using consistent exposure settings for FITC and Alexa Fluor® 647. For quantification of CD44 expression and the mannosylation score using fluorescent images, background fluorescence was subtracted uniformly across all images. Using Zen2 software (Zeiss core imaging software) ROIs were drawn around each tissue core, excluding any folded or damaged areas. Mean fluorescence intensity values for GNL-FITC (mannose) and Alexa Fluor® 647 (CD44) were measured and normalized onto a 1–100 scale by assigning the lowest measured intensity a value of 1 and the highest a value of 100. These normalized data were then used to determine the correlation between CD44 expression and mannosylation score.

### Knockdown of LMAN1 and LMAN2

For M1123 cells, the mannose-binding lectins LMAN1 and LMAN2 were knocked down using liposomal transfection with LMAN 1/2 siRNA (Millipore-Sigma, Cat# SASI_Hs01_00189318 and SASI_Hs01_00209153), and LMAN1/2 expression was quantified with qRT-PCR and normalized to GAPDH. siRNA transfections were performed using RNAiMax (Thermo Fisher Scientific) according to the manufacturer’s recommendations. Briefly, GBM neurosphere single-cell suspensions (2.5×105 cells) were seeded onto ultra-low-attachment 24-well plates (Millipore-Sigma) in 800 μl of neurosphere medium. Lipofectamine RNAiMax reagent (12 μl) was added to 100 μl of Opti-MEM medium. In a separate tube, siRNA oligos were diluted in 100 μl of Opti-MEM to achieve a final concentration of 240 nM. Tubes were combined, mixed and incubated at RT for 15 minutes to allow lipid-siRNA complex formation. The mixture was then added dropwise to GBM neurospheres single-cell suspensions. Cells were collected 5 days after transfections for gene expression analysis.

### qRT-PCR

Total RNA was extracted from the cells using a RNeasy mini kit (Qi-agen, Germantown, MD.cDNA was made by reverse-transcribing 1 μg of total RNA using MuLV Reverse Transcriptase and Oligo (dT) primers (Applied Biosystems,Waltham, MA). qRT-PCR was performed with the Bio-Rad CFX detection System (BioRad, Hercules, CA), and the expression of target genes was measured using the Power SYBR green PCR kit (Applied Biosystems). The samples were amplified in triplicate, and relative gene expression was analyzed using Bio-Rad CFX manager software v3.1 and normalized to 18S RNA. The primer sequences used in this study were obtained from PrimerBank (https://pga.mgh.harvard.edu/primerbank/, last accessed on 12.20.24) and are listed in **Table S1**.

### *In vitro* MANw CEST MRI

For *in vitro* MANw CEST MRI studies, 1×10^6^ GBM1a and M1123 neurospheres with an average size of 500 µm were collected, rinsed three times with PBS, and loaded into 5 mm NMR tubes (Wilmad®, USA). CEST MRI was then conducted on an 11.7 T Bruker Biospin vertical bore scanner equipped with a 20 mm birdcage transmit/receive coil. A modified RARE (rapid acquisition with refocused echoes) sequence was employed, with the following parameters: repetition time (TR)/echo time (TE)=6,000/5 ms, RARE factor=32, number of averages (NA)=2, slice thickness=1 mm, field of view (FOV)=16×16 mm, matrix size = 64×64, in-plane resolution=0.25×0.25 mm, B_1_=2.4 µT, and saturation time (T_sat_)=4 s. Saturation frequencies ranged from −5 to +5 ppm in 0.2 ppm intervals, with 0 ppm referencing the water resonance. The total acquisition time wass 20 min and 48 s. LMAN1-, LMAN2-, and dual LMAN1/2–knockdown neurospheres were subsequently imaged under identical conditions and compared with wt M1123. All experiments were performed as three independent replicates.

### Animal Model

All animal procedures were approved by the Johns Hopkins University Animal Care and Use Committee. Immunodeficient NSG mice (NOD.Cg-Prkdc^scid^ Il2rg^tm1Wjl^/SzJ) were obtained from Jackson Laboratories (Bar Harbor, ME, USA) and maintained as an in-house breeding colony. Male and female mice, aged 8 weeks (22–25 g), were housed under a 12-hour light/dark cycle with unrestricted access to food and water. During all surgery and imaging procedures, mice were anesthetized with isoflurane (2–3% for induction and 1–2% for maintenance in O_2_), using a calibrated vaporizer and scavenging system. Breathing rate and body temperature were continuously monitored and controlled throughout the scanning procedure. For stereotaxic injections, a burr hole was drilled at coordinates 0 mm caudal and 2 mm lateral to the bregma on each side. A Hamilton syringe was used to inject 2.5 µL of single cell suspensions from neurospheres. Neurospheres were dissociated into single cells using gentle mechanical trituration and enzymatic treatment with Accutase. Spheres were first pipetted gently using a wide-bore tip, followed by incubation in pre-warmed StemPro™ Accutase™ cell dissociation reagent (Gibco, USA) at 37°C for 20 minutes, with occasional swirling to obtain a single-cell suspension. Cells (1.5×10^5^ M1123 or GBM1A in PBS) were injected into the right and left striatum at a depth of 2 mm below the endocranium, administered at a rate of 0.2 µL min^−1^. To further verify that the CEST signal is generated by mannose in mesenchymal cells, LMAN1/2 KD neurospheres were injected into the left striatum and wild-type M1123 neurospheres into the right striatum using the same procedure in three mice. To compare the perfusion properties of GBM1a and M1123 tumors, mice received 0.2 mmol/kg of a Gd-based contrast agent (Omniscan, Nycomed, Oslo, Norway) via a pre-inserted intravenous catheter. Dynamic T1-weighted imaging was conducted before injection (baseline) and continuously for 10 minutes post-injection to evaluate perfusion and vascular permeability.

### *In vivo* ManW CEST, APTw, and Gd-enhanced T1w MRI

MRI was carried out on days 1, 8, and 16 post-transplantation using an 11.7 T Bruker Biospec preclinical scanner (Bruker, Ettlingen, Germany) with a 23-mm volume transmit/receive coil coil. T2-weighted (T2w) MRI parameters were TR/TE=3,000/5.6 ms, RARE factor=16, slice thickness=1 mm, FOV=1.6×1.5 cm, matrix=256×128, and NA=3. MANw CEST and APTw MRI parameters were TR/TE=5,500/3.7 ms, RARE factor=23, single slice, slice thickness= 1 mm, FOV=1.6×1.6 cm, matrix=64×64, NA=1, B_1_=2.4 µT, and T_sat_=3 s, with saturation frequencies ranging from −5 to +5 ppm in 0.25 ppm increments (water signal set to 0 ppm). The total acquisition time was 7 minutes. Tumor and brain regions of interest (ROIs) were manually drawn from T2-w images. This experiment was performed in five independent replicates. Gd-enhanced MRI was performed by i.v. injection of ProHance® (0.1 mmol/kg, bolus over about 10 s, injection volume =50 µL, injection rate= 5 µL/s). A single-slice FLASH gradient echo sequence (TR/TE=18/3 ms, and flip angle =15°) was used to acquire DCE images at the same position and spatial resolution as the CEST images. The acquisition started two minutes before injection. The total acquisition time of DCE MRI was 13 min at temporal resolution of 14.6 s.

### Post-mortem tissue analysis

Mice were transcardially perfused with 10 mM PBS and 4% paraformaldehyde (PFA). The brain was removed and re-immersed in 4% PFA at 4 °C for 24 h, then transferred to 30% sucrose for another 72 h. After embedding in optimal cutting temperature compound (OTC), the brain was cryosectioned into 10 μm sections. For GNL and anti-CD44 staining, brain sections were first rehydrated with PBS for 10 min and then incubated with 20 μg/ml FITC-GNL and 1:250 Alexa Fluor® 647 anti-CD44 antibody for 1h at RT. After washing 3x with PBS, sections were coverslipped using mounting medium containing DAPI. Fluorescence microscopy was performed using a Zeiss Axiovert 200 M inverted epifluorescence microscope.

### Statistics and reproducibility

For statistical analysis of patient data, pairwise comparisons between group levels with corrections for multiple testing (p values with Bonferroni correction) were used. Pearson’s correlation analysis was performed to assess relationships between CD44 expression and mannosylation score, and linear regression models were applied to visualize trends with 95% confidence intervals. Data are presented as mean±standard deviation (SD and were analyzed using a one-way analysis of variance (ANOVA), followed by Dunnett’s post hoc test for multiple group comparisons, with significance thresholds set at *p<0.05, ** p<0.01, and ***p<0.001 and ****p<0.0001. Comparisons between tumors were performed using a two-tailed Student’s t-test. Sample sizes were determined to achieve adequate statistical power (>90%, p=0.01) for detecting expected effect sizes, estimated based on our preliminary data and prior experience with similar experiments. The number of independent repetitions for each experiment is given in the corresponding figure captions. All statistical analyses were conducted using GraphPad Prism 6.0. Investigators remained blinded to group allocations during and after experiments including MRI data processing.

## Acknowledgements

This study was funded by NIH grants R01 EB030376 (JWMB), R01 NS120949 (HLB), R01 NS073611 (JL), R01 NS096754 (JL), R01 CA261974 (GL), the Maryland Stem Cell Research Fund 2023-MSCRFD-6135 (JWMB), the Sidney Kimmel Comprehensive Cancer Center Translational Research Central Services Shared Resource (P30CA006973), and the Clinical Translation Core at the Kennedy Krieger Institute (5P50 HD 103538). We thank Bachchu Lal for performing part of the cryosectioning, and Dr. Calixto-Hope G. Lucas Jr. for his advice on the glioblastoma tissue array and neuropathology.

## Author contributions

B.G. and J.W.M.B. conceived the ideas for initiating this study. H.L-B. performed computational analysis of transcriptomic datasets. B.G., H.L-B., G.L., and J.W.M.B. designed the methods and experimental design. B.G., H.L-B., S.K. S.S, C-H.G.L.Jr. and G.L. performed *in vitro* cell and *in vivo* animal experiments and data analysis. B.G. and H.L-B. wrote the initial manuscript draft. J.W.M.B. and H.L-B. acquired the funding for this project. All authors read, edited and revised the manuscript.

## Inclusion & Ethics Statement

All co-authors have made substantial contributions to the study’s conception, design, data collection, and interpretation. Each has reviewed and approved the final manuscript. This research complied with all relevant ethical guidelines and received appropriate institutional approvals, where required. No conflicts of interest are declared.

## Supplementary Information

**Table S1:**
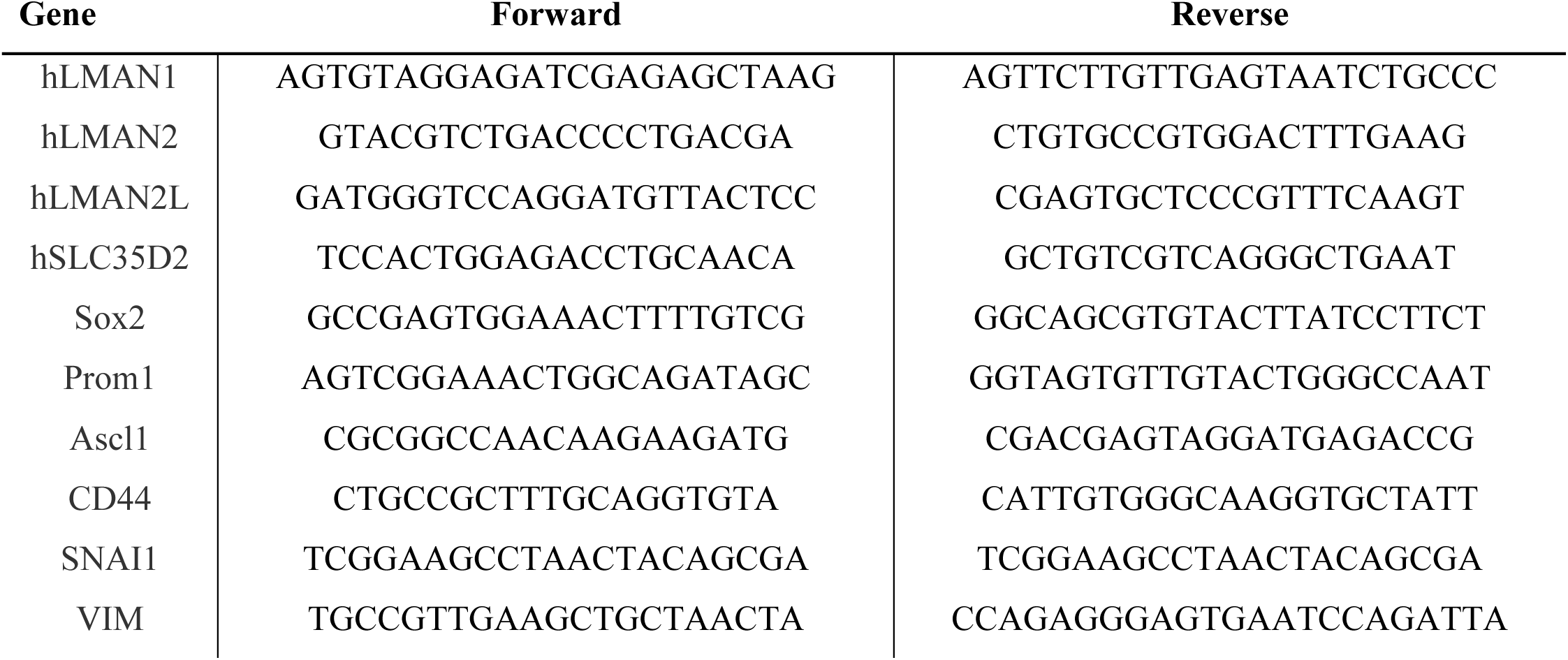
Primer sequences used for qRT-PCR analysis.

**Figure S1:**
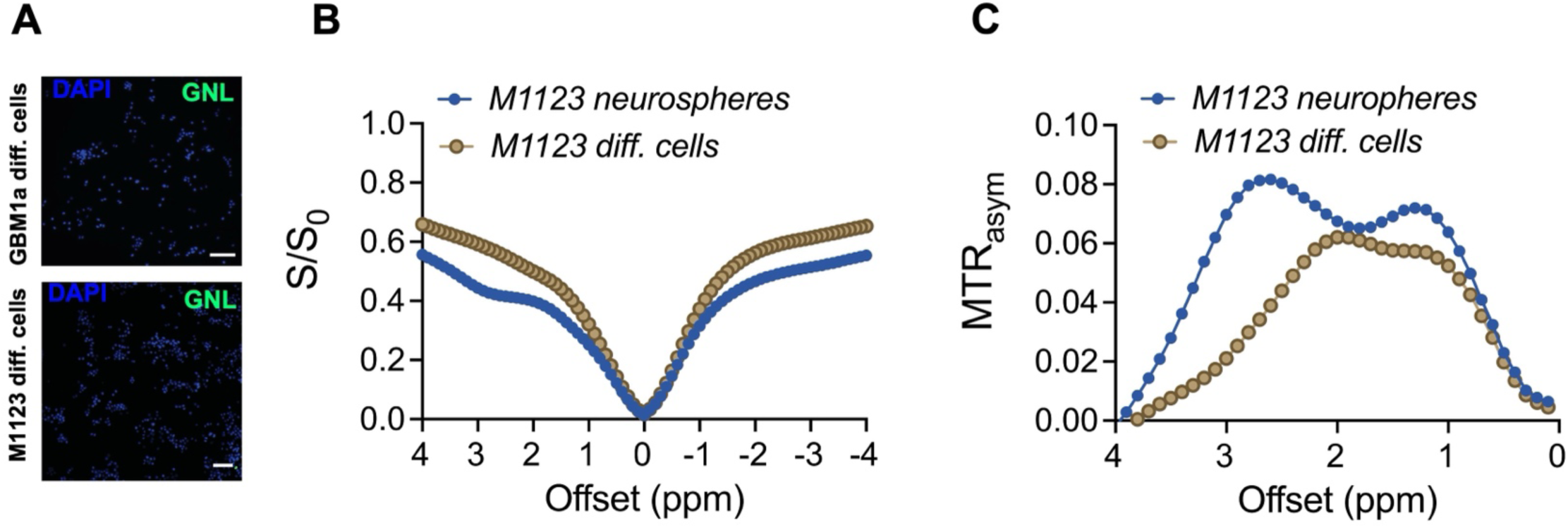
(**A**) *In vitro* GNL-FITC staining for mannosylation. (**B,C**) *In vitro* Z-spectra (**B**) and (**C**) ΔMTR_asym_ spectra of serum-induced differentiated and M1123 neurospheres. Scale bar in A=100 μm.

**Figure S2:**
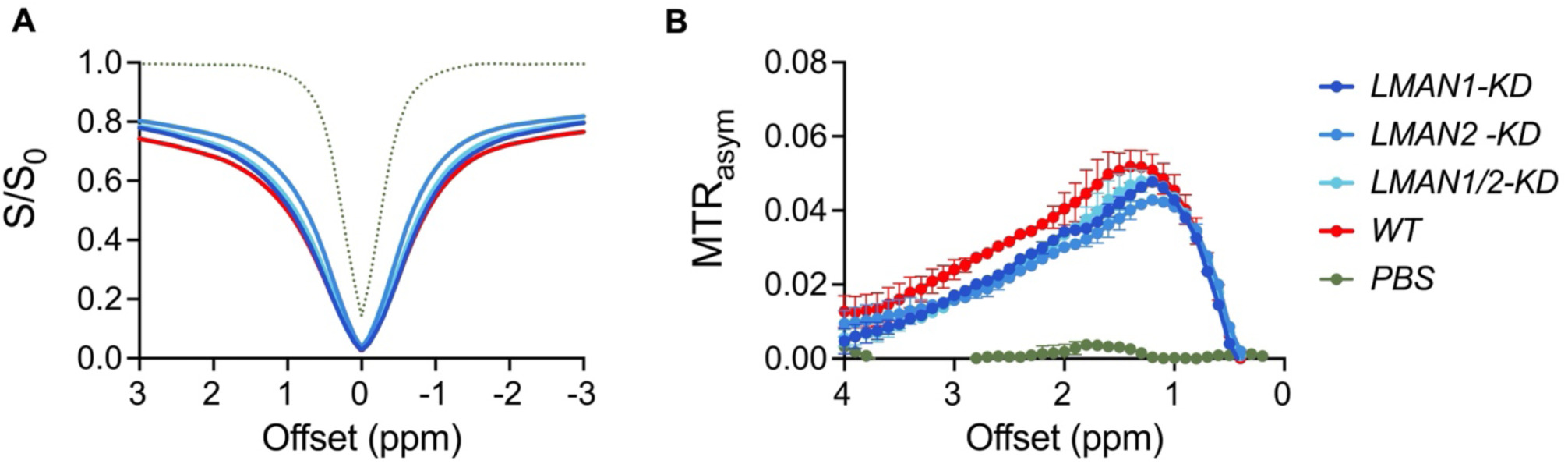
(**A**) *In vitro* Z-spectra and (**B**) ΔMTRasym spectra of LMAN1, LMAN2, and LMAN1/LMAN2 knocked down (KD) and wild type (WT) M1123 neurospheres.

**Figure S3:**
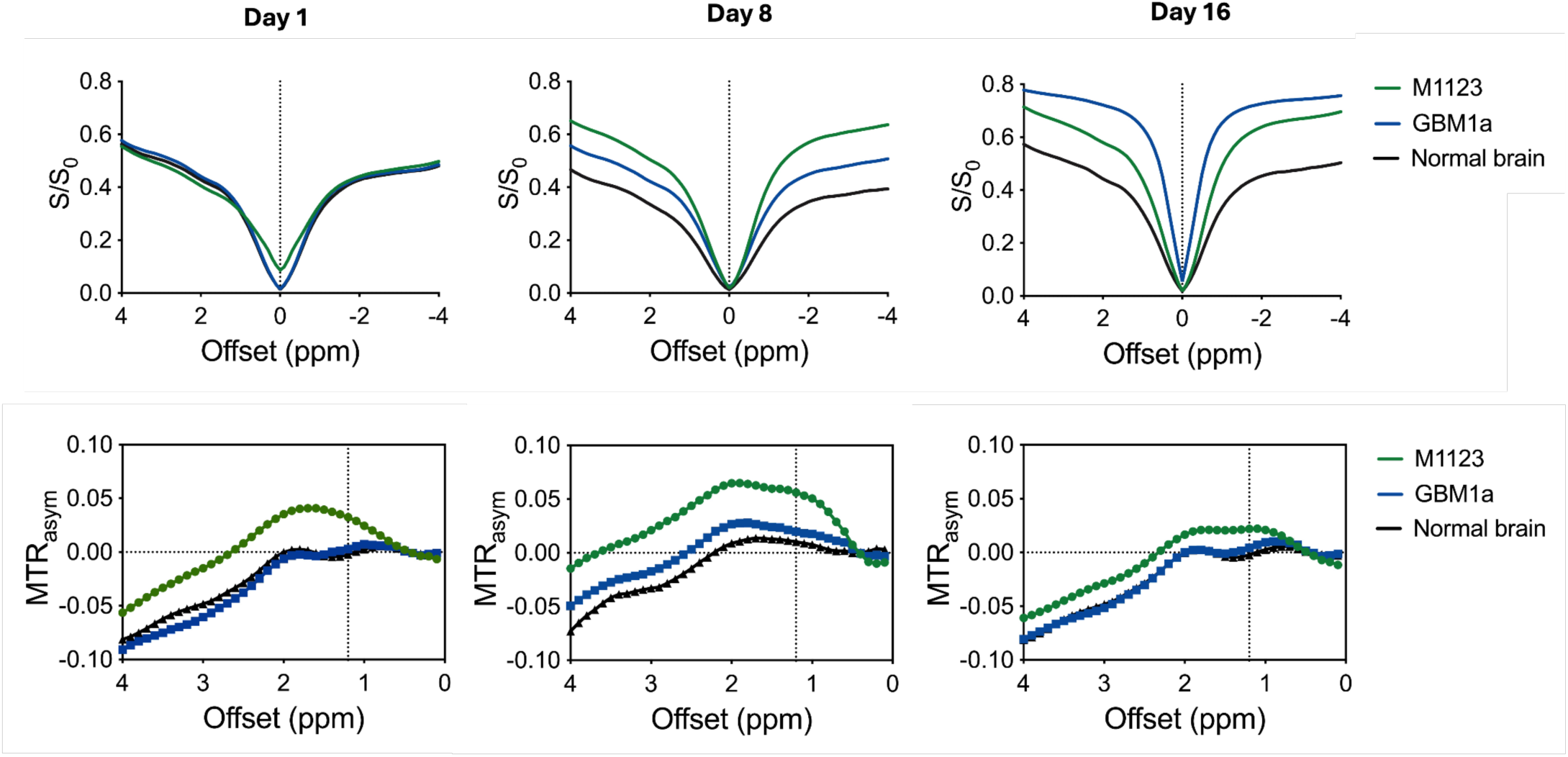
Z-spectra (top row) and ΔMTR_asym_ spectra (bottom row) of GBM1a and M1123 cells implanted bilaterally in mouse brain for the ROIs drawn on day 1, 8 and 16 post inoculation (n=5 animals). Vertical dashed lines are set at 1.2 ppm at which saturation frequency the in vivo MANw CEST MR images were obtained. Note that the relative widths of Z-spectra vary at different time points. This variation occurs since Z-spectra widths are dependent on the T1 relaxation times, where a narrower shape corresponds to a longer T1GBM1a showed a marked T1 increase between day 8 and 16.

**Figure S4:**
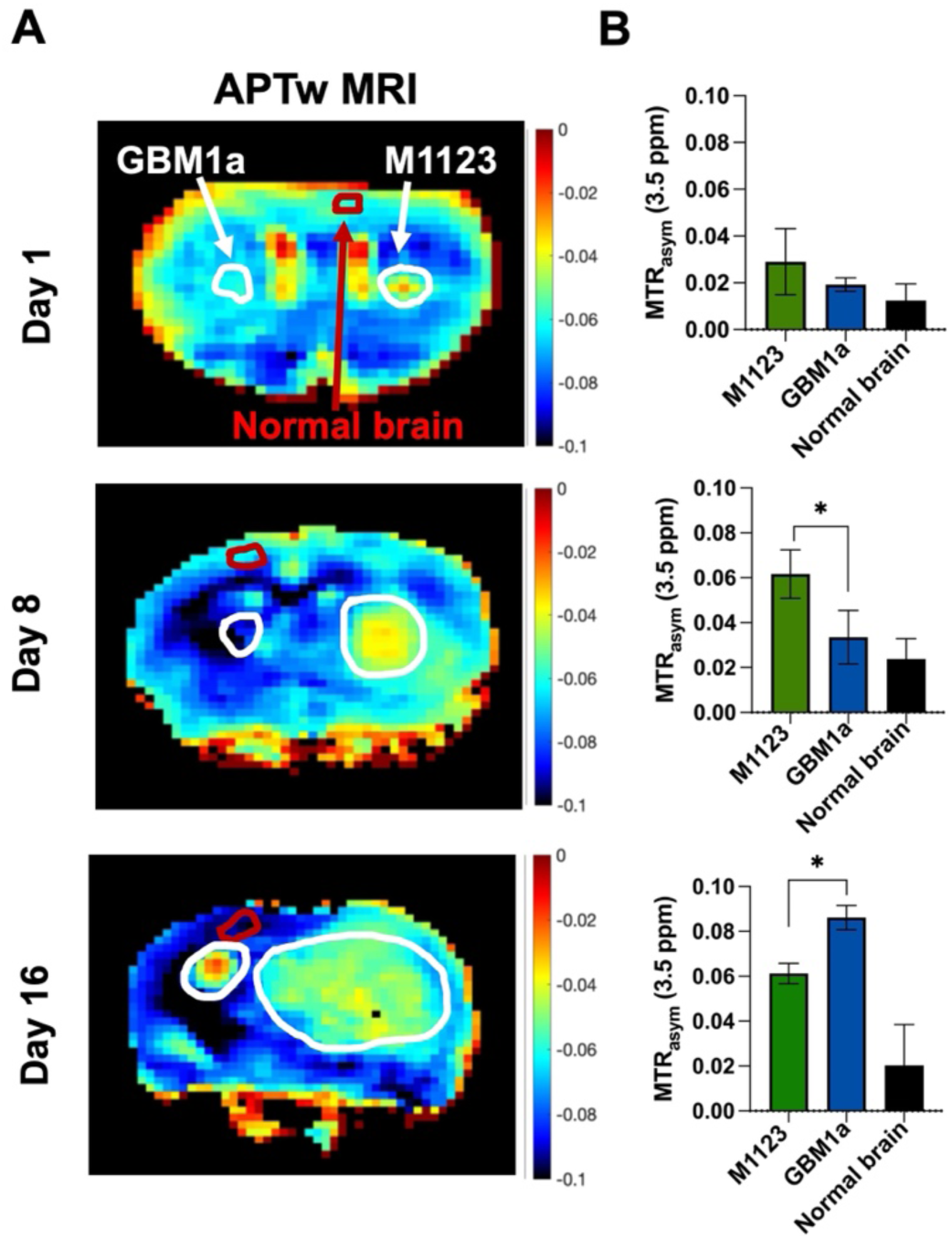
(**A**) Serial APTw MRI at different time points post-tumor implantation. (**B**) Quantified APT CEST MRI signal at 3.5 ppm for the regions of interest as a representative example for n=3 animals.

**Figure S5:**
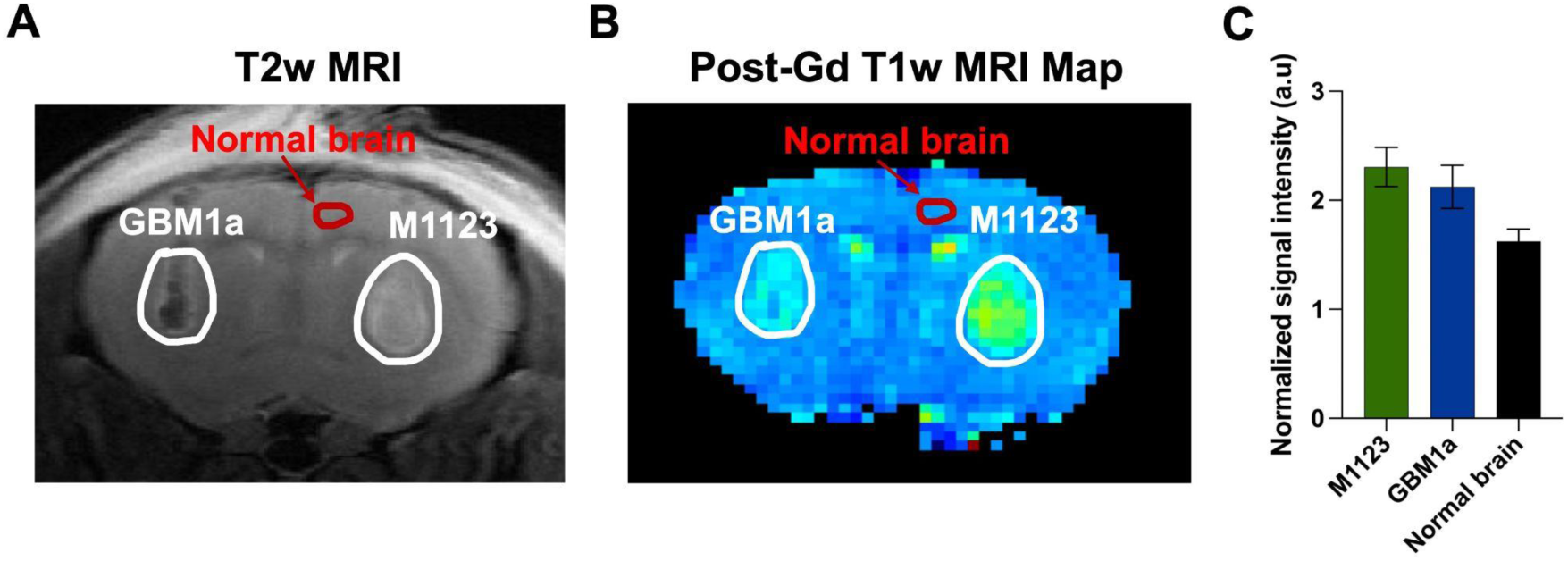
(**A**) T2-weighted and (**B**) Gd-DTPA enhanced T1-weighted map of GBM1a and M1123 implanted bilaterally in mouse brain, 8 days post-inoculation. (**C**) Quantified signal intensity of Gd-enhanced T1-weighted MR images for M1123 and GBM1 compared to normal brain.

